# An *in vitro* model of acute horizontal basal cell activation reveals dynamic gene regulatory networks underlying the acute activation phase

**DOI:** 10.1101/2023.12.14.568855

**Authors:** Camila M. Barrios-Camacho, Matthew J. Zunitch, Jonathan D. Louie, Woochan Jang, James E. Schwob

## Abstract

Horizontal basal cells (HBCs) activate only in response to severe olfactory epithelium (OE) injury. This activation is mediated by the loss of the transcription factor TP63. Using the compound phorbol 12-myristate 13-acetate (PMA), we find that we can model the process of acute HBC activation. First, we find that PMA treatment induces a rapid loss in TP63 protein and induces the expression of HOPX and the nuclear translocation of RELA, previously identified to mediate HBC activation. Using bulk RNA sequencing, we find that PMA-treated HBCs pass through various stages of acute activation identifiable by transcriptional regulatory signatures that mimic stages identified *in vivo*. These temporal stages are associated with varying degrees of engraftment and differentiation potential in transplantation assays. Together, this data shows that our model can model physiologically relevant features of HBC activation and identifies new candidates for mechanistic testing.

## Introduction

The only neurons in the human body that directly interface the external environment are housed in the olfactory epithelium (OE), a specialized chemosensory neuroepithelium lining a portion of the nasal cavity(Carr et al., 2001). In response to continuous exposure and injury, the OE has evolved a remarkable and nearly life-long capacity to regenerate its neuronal and non-neuronal constituents(Schwob, 2005; Schwob et al., 2017). The extent of the regeneration makes it unique by comparison to the other few sites in the nervous system where adult neurogenesis takes place. Crucial to this process is the activity of two stem cell populations: the constitutively proliferating and differentiating globose basal cells (GBCs), and the horizontal basal cells (HBCs), which only activate in response to severe epithelial injury. Both populations possess multipotent capacity within the context of epithelial regeneration(Carter et al., 2004; Jang et al., 2003). In the anosmic, aged olfactory epithelium, focal sites of aneuronal OE and respiratory metaplasia can be observed. In both states, the GBC population is lost, while HBC population fails to activate and regenerate the neuronal compartment. Consequently, understanding the cohort of regulators that facilitate the HBC transition out of dormancy post-lesion would facilitate the clinical use of HBCs for the treatment of anosmia.

Previous studies have shown that the loss of the transcription factor *Tp63*, a member of the *Tp53* family of transcription factors, is necessary and sufficient to release of HBCs from dormancy and into active multipotency *in vivo*(Schnittke et al., 2015; Schwob, 2005; Schwob et al., 2017). On the other hand, viral delivery of *Tp63* into activated HBCs is sufficient to return them to dormancy(Schnittke et al., 2015). TP63 loss and the subsequent activation of *HBCs* can be achieved in vivo with selective ablation of the sustentacular (Sus) cell population, whereas selective ablation of the neuronal population via olfactory bulbectomy while lowering the activation threshold is insufficient to release HBCs from dormancy on its own(Herrick et al., 2017). Similarly, Notch1 knockout lowers the threshold for HBC activation in the uninjured state thus implicating Notch signaling in the maintenance of HBC dormancy(Herrick et al., 2017). Additionally, NF-kB signaling has been shown to be a crucial effector of HBC activation as HBC-specific deletion of the NF-kB transcriptional effector *Rela* blunts regeneration of the OE after methimazole lesion(Chen et al., 2017). Lastly, while HBCs can return to dormancy following injury, expression of *Hopx* is selectively enriched in differentiation-committed HBCs and can be used as a marker for activation (Gadye et al., 2017).

Previous transcriptomic analyses of HBCs both at the bulk and single-cell level have highlighted major branching points underpinning HBC diversity and lineage specification in the post-activation setting. After lesion to the OE, HBCs exhibit divergent differentiation into either Sus cells or GBCs, the latter of which may then differentiate into neurons, duct/glad cells (D/G), microvillar (MV) cells, or additional Sus cells(R. B. Fletcher et al., 2017; Gadye et al., 2017). The molecular mechanisms of underlying these HBCs fate determination steps are unknown.

A unifying limitation in understanding HBC activation is poor temporal resolution and the challenges of *in vivo* manipulation. While lineage-tracing studies have been performed on injury-activated HBCs *in vivo*, these studies focused on the events occurring twenty-four hours or more after lesion(Gadye et al., 2017). While previous studies from our lab found that sustained treatment of cultured HBCs with retinoic acid (RA) eventually induces the loss of *Trp63,* treatment times were extensive (3+ days)(Peterson et al., 2019). Thus, there is an urgent need to develop *in vitro* systems that enable robust, timely, and physiologic activation of HBCs to better study the molecular regulation of acute activation, HBC heterogeneity, and fate determinism in the acute activation period.

To that end, we present a new *in vitro* platform for achieving physiologic and reversible HBC activation within twelve hours. This model recapitulates many aspects of the acute molecular and developmental processes that govern HBC-mediated regeneration of the olfactory epithelium. Using this model, we analyzed loss of the master dormancy regulator TP63 with fine temporal resolution, identifying the precise window during which TP63 undergoes precipitous decline. Additionally, we find a transient and robust nuclear translocation of the NF-κB effector RELA preceeding precipitous TP63 loss. This activation model unlocks engraftment potential and generates a heterogenous HBC population that can self-renew or differentiate into GBCs, in a manner consistent with post-lesion behavior of HBCs *in vivo*. Finally, we study the transcriptional regulatory signatures throughout the activation process and identify regulator candidates that may be responsible for early fate-determining events in acutely activated HBCs.

## Results

While previous analyses of *in vivo* lesioned HBCs have revealed near-absent expression of TP63 at 18 hours post methyl bromide (MeBr)(Schnittke et al., 2015), the kinetics of acute TP63 loss following injury is unknown but critical to the evaluation of any *in vitro* model of injury-induced activation. To map the *in situ* kinetics of TP63 loss during the acute phase of OE lesion, we performed a time course of methimazole (Mtz)-induced injury in wild type mice. To do so, we used confocal microscopy paired with a high-throughput CellProfiler-mediated image analysis to quantify nuclear TP63 immunofluorescence levels over time in 19,494 HBCs(McQuin et al., 2018). TP63 levels were assessed on a per-cell basis at 0, 6, 12, 24, and 48 hours post MTZ injection (hours post injection; HPI). We also allowed the tissue to partially regenerate and quantified TP63 expression at 7 days post injection (DPI), after a resting HBC population is re-established (Figure 1A). We observed a high degree of HBC heterogeneity during the acute activation phase, with some HBCs demonstrating rapid responsiveness to injury (indicated by rapid TP63 loss) and others demonstrating a degree of resistance (preservation of near-baseline TP63 expression or minimal decline) (Figure 1B). This observed heterogeneity prompted us to classify our HBCs into three populations, utilizing TP63 immunofluorescence labeling as the principal determinant: a resting population, a transitioning population, and an activated population (Figure 1C). Results from this analysis suggest that multiple populations of HBCs exist in all conditions of the OE, including unlesioned tissue. Injury induces compositional changes to the HBC population; at 12 HPI, 95% of HBCs are in an activated or transitioning state, while retaining a small (5%) dormant fraction (see Supplemental Information S1). Lastly, analysis of recovering tissue (7DPI) indicates that HBC composition has not returned to the unlesioned context, despite a significant reestablishment of the resting population (Time Point 0 = 74.9% resting, 14.3% transitioning, 10.6% activated vs. Time Point 168 HPT = 51.6% resting, 25.1% transitioning, 23.1% activated). Together, these data show that HBC activation is not a synchronous process, and that multi-stage HBCs compose the OE across all tissue contexts. Most importantly, we observe that overall TP63 expression *in vivo* reaches *a nadir* at 12 hours post activation.

**Figure 1.**
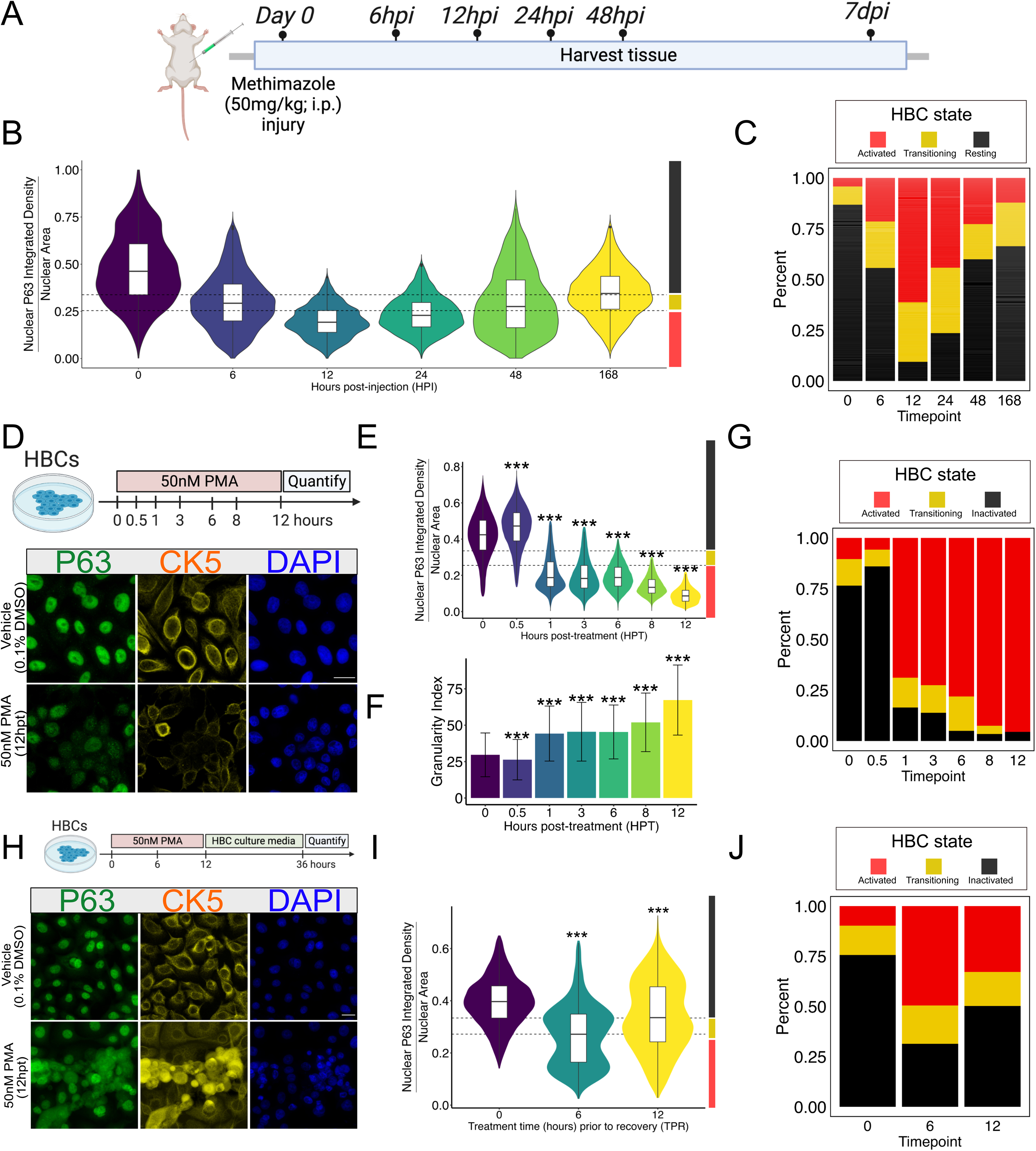
PMA treatment leads to TP63 degradation *in vitro* but is not deterministic. A) Schematic illustration of MTZ injury timecourse. B) An integrated violin and boxplot representing the Nuclear P63 Integrated Density/Nuclear Area quantification results of each time point (n=2). C) Bar plot showing proportional composition of HBC state at each time point. D) Schematic illustration of the PMA-mediated HBC activation assay (top). Representative sum projected z-stack confocal 63x images of vehicle-treated and 12h-long 50nm-treated HBCs (below). E) An integrated violin and boxplot representing the Nuclear P63 Integrated Density/Nuclear Area quantification results of each time point (n=3). F) An integrated violin and boxplot representing the Granularity I output from CellProfiler. G) Bar plot showing proportional composition of HBC state at each timepoint post-PMA treatment. H) Schematic illustration of the PMA-mediated HBC recovery assay (top). Representative sum projected z-stack confocal 63x images of vehicle-treated and 12h-long 50nm-treated HBCs (bottom). I) An integrated violin and boxplot representing the Nuclear P63 Integrated Density/Nuclear Area quantification results of each time point (n=3). J) Bar plot showing proportional composition of HBC state at each time point post-PMA treatment and recovery. For (B, E, I). Dashed lines represent TP63 fluorescence expression level thresholds corresponding to HBC state. Right y-axis color blocks on (B,E,I) correspond to these states, shown in (C,G,J). Extreme outliers were removed utilizing the Tukey method (1.5 x IQR) (E, I). For (E,F,I) statistical significance was determined by one-way ANOVA followed by Dunnett’s test. For (E, F, I): *p< 0.05,**p< 0.001,***p< 0.0001 (E,I). For (F), mean and standard deviation is reported. Scale bars: 20 μm (D,H). See Supplemental Information S1 for information about sample sizes and number of cells per condition.

### Treatment with PMA induces activation *in vitro* through the rapid loss of TP63

We set out to develop an *in vitro* assay capable of modeling and probing the underlying molecular mechanisms of acute HBC activation (Figure 1D, top). To induce activation, we utilized phorbol 12-myrsitate 13-acetate (PMA), which is an activator of numerous signaling pathways previously implicated in HBC activation, such as the NF-κB pathway(Holden et al., 2008). Following treatment of *in vitro* HBCs with PMA, we assayed TP63 expression in a manner like the *in vivo* analysis at 6 time points over a 12-hour period to assess the kinetics and extent of TP63 loss. Given that injury-mediated TP63 loss *in vivo* is not homogenous nor linear, we prioritized the ability to quantify levels of TP63 at a single-cell resolution. Results from this experiment show a triphasic change in TP63 immunofluorescence over the 12-hour time course. Thirty minutes after PMA treatment, we observed a ∼10% increase of TP63 protein compared to vehicle, possibly reflecting homeostatic mechanisms protecting TP63 pools (Figure 1E). However, levels rapidly fall between 30 minutes and one-hour post-treatment to approximately ∼52% of vehicle-treated cells. Levels then appeared to stabilize between 1 and 6 HPT before undergoing a second phase of reduction between 6 and 12 HPT. At the end of the 12-hour treatment period, HBCs lost ∼78% of their nuclear TP63 immunofluorescence. Visual inspection of high magnification images (Figure 1D, bottom) shows the emergence of a granular, punctate staining pattern of TP63 in both the nucleus and cytoplasm, likely representing sites of TP63 sequestration and degradation. This change in immunofluorescence texture was quantified using the MeasureGranularity module in CellProfiler, which revealed linearly increasing levels of punctate TP63 protein (Figure 1F). While most cells rapidly responded to treatment, a minor fraction of HBCs at 8 and 12 HPT maintained near-baseline TP63 immunofluorescence levels and thus appeared resistant to PMA treatment (Figure 1G).

In the acute post-injury environment, the HBC population bifurcates into cells that are fated to differentiate versus those that re-establish dormancy. Having established that PMA induces the loss of TP63, we then sought to assess whether this physiologic plasticity was maintained in the PMA-activated HBCs *in vitro.* We assessed this by testing whether they are capable of re-establishing baseline TP63 levels after a transient PMA activation. To test this, we treated HBCs with PMA for either 6 or 12 h before washing out the drug and allowing cells to recover in maintenance media for 36 h (Figure 1H, top). Nuclear TP63 immunofluorescence was then analyzed as above with CellProfiler. Results from this experiment showed that HBCs can resynthesize TP63 following acute PMA treatment, with HBCs experiencing a net loss of only 13% of TP63 protein after 12h of PMA treatment and recovery (Figure 1I). Interestingly, we observed a clear bifurcation in the population when cells were treated with PMA for 6 hours and then transitioned back to maintenance media. Under this paradigm, roughly half of the cells re-established vehicle-treated TP63 levels whereas the other half continued to lose TP63 in a manner consistent with activation (Figure 1J). Additionally, we observed a three-dimensional reorganization in the culture. Whereas vehicle-treated HBCs form a stereotyped cobblestone monolayer, cultures that were transiently activated contained regions of piled-up HBCs (Figure 1H, bottom). Together, these results suggest that a transient activation event in this system may be leveraged to study physiologic subpopulations of HBCs with characteristics that mirror those in the acute post-injury setting *in vivo*.

### Bulk RNA-sequencing of PMA-treated HBCs reveals transcriptional networks important for HBC activation

Having established a model system for the early stages of HBC activation, we assessed transcriptomic changes during this process using bulk-RNA sequencing of vehicle-treated HBCs and two populations of PMA-treated HBCs (6 HPT and 12 HPT), which henceforth will be referred to as acute-activation and extended-activation HBCs, respectively. Results from principal component analysis indicate that 92% of the variance in the data is explained by whether the cells were treated with PMA and 6% of the variance can be explained by the length of PMA treatment time (6h vs 12h), providing us the opportunity to analyze subtle, but potentially important changes in the transcriptome (Supplemental Figure 1A). Analysis of differentially expressed genes (DEGs; Treatment/Vehicle) in the PMA-treated conditions revealed a core PMA-induced treatment signature, irrespective of length of treatment; both time points shared 6458 DEGs (Supplemental Figure 1B).

Furthermore, of these 6458 DEGs, 96% were regulated in the same direction as compared to vehicle (e.g., upregulated or downregulated), showing that PMA treatment has a persistent effect on the DEGs at both time points. We next evaluated the gene ontology networks that were either up- or down-regulated in our 6 HPT/Vehicle and 12 HPT/Vehicle specific signatures (6 HPT = 1388 genes; 12 HPT = 1271 genes) as well as the gene ontology networks induced by PMA treatment alone (shared DEGs: 6458). We utilized Metascape to identify significantly enriched pathways across these six contexts(Zhou et al., 2019). At 6 HPT, the ontology networks transcriptional machinery, NF-κB signaling, and mRNA modifications are upregulated and enriched (Supplemental Figure 1C, D). In contrast, this analysis revealed that networks for membrane trafficking, mitochondrial transport, and the regulation of cell-matrix adhesion were down-regulated (Supplemental Figure 1C, D). Results from 12 HPT analysis reveal upregulated MAP2K signaling, cytoskeletal remodeling, and axon development (Supplemental Figure 1E, F). In contrast, we found downregulated signatures involving cell cycle, ncRNA processing, and RNA polymerase III transcription (Supplemental Figure 1E, F). In addition to the time-point specific ontology analysis, we also interrogated the ontology networks specifically induced by PMA, irrespective of length of treatment (Supplemental Figure 1G,H). In sum, we found that treating HBCs with PMA upregulated various morphogenic, developmental, and differentiation ontology categories, while simultaneously downregulating cell cycle, DNA replication, and cell division categories. These results, paired with the time-point specific ontology changes, show that distinct metabolic and signaling pathways are recruited as the transcriptome shifts across the activation landscape.

### PMA treatment of cultured HBCs induces *in vivo* activation genes

We next compared the gene expression of HBCs activated via PMA *in vitro* and HBCs activating natively under injury conditions *in vivo*. Given the potential wide-ranging effects of PMA on cultured cells, we assessed transcriptomic similarity using specific established marker genes and by analyzing the collective expression of entire gene sets. For our *in vivo* reference, we re-analyzed a single cell data set of resting and activated HBCs from the wild-type mouse OE(Gadye et al., 2017). We clustered Krt5(+) HBCs into three groups dependent on their gene expression of dormancy regulator *Trp63,* proliferation marker *Mki67,* and *bona fide* activation marker *Hopx.* This dataset consisted of the following three distinct HBC populations, in developmental order. First, we identified a resting state defined by cells that have high *Trp63* expression, negative for the proliferation marker *Mki67*, and negative for the activation marker *Hopx*. Second, we identify a partially activated, proliferating state, demarcated by medium *Trp63* expression, positive *Mki6*7 expression, and negative *Hopx* expression. Lastly, we identify an activated population, indicated by low-to-absent *Trp63* expression, negative *Mki67*, and positive *Hopx* expression. (Figure 2A, 2B). We then used these population to map gene expression occurring *in vitro* back to native cell types. We additionally immunostained our PMA-activated HBCs for HOPX protein, finding an upregulation of HOPX at 12 HPT, indicating that PMA-treatment can induce this specific *in vivo* HBC state (Figure 2C).

**Figure 2.**
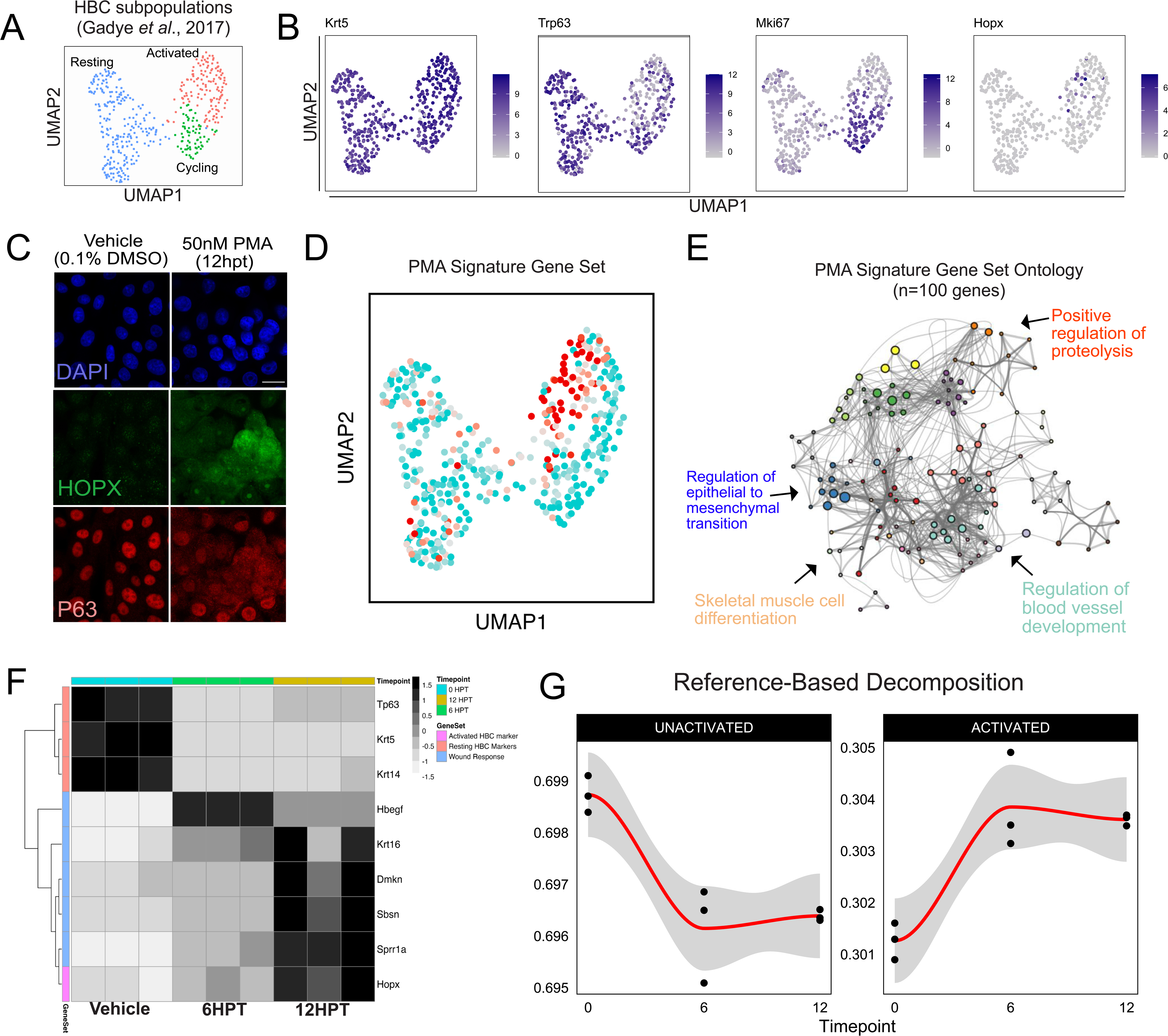
PMA-treated HBCs correspond to *in vivo* HBCs state. A) Clustering of HBC subpopulations found in the regenerating olfactory epithelium (Gadye et al. 2017) B) Expression of marker genes differentiating the HBC subpopulations: pan-HBC markers -*Krt5*; cycling-HBC marker – *Mki67*; dormant-HBC marker - *Trp63*; activated HBC – *Hopx* C) Representative immunostaining for HOPX, an activated HBC marker previously identified. Cells were treated for 12 h with PMA before being subjected to staining. Scale bar: 20 μm. D) Relative signature expression in HBCs. Quantity reflects the collective expression of the gene set relative to a randomized control set of the same size (see Tirosh *et al*, 2016.). The “PMA signature” is comprised of the top 100 differentially upregulated genes in response to PMA-induced activation *in vitro*. Only *Sprr1a* is common between the gene sets. Cyan indicates low signature expression; red indicates high signature expression. E) Gene ontology network visualizing and summarizing the Biological Process terms that are enriched in the 100-gene PMA signature. F) Heatmap visualizing the expression of HBC markers and wound response genes detected in the PMA-treated *in vitro* HBCs. G) Reference-based deconvolution of the bulk RNAseq samples based on the single cell gene expression patterns of inactivated and activated HBCs *in vivo*. This approach relies on the raw single cell data to estimate absolute cell type proportions.

Using the module detection function of Seurat (Tirosh et al., 2016), we then plotted the collective expression of the top 100 differentially expressed genes induced by PMA treatment, finding elevated expression of this signature in the same activated HBC cluster (Figure 2D). Importantly, only one gene of the hundred-gene PMA signature overlapped with the wound response gene set (*Sprr1a*) indicating that PMA-induced genes, distinct from wound response genes, are indeed upregulated during *in vivo* activation. Gene network ontology of these same genes indicated enrichment for signaling, developmental, and differentiation associated categories (Figure 2E).

We then analyzed the bulk RNAseq data of the *in vitro* HBCs to determine whether the transcriptomic changes induced by PMA activation recapitulated known gene expression changes in the post-lesion environment. Looking specifically at resting HBC markers and previously characterized activation-associated wound response genes (Gadye et al., 2017) and *Hopx*, we observed that PMA treatment caused a decrease in *Tp63, Krt5*, and *Krt14* expression concurrent with unanimous upregulation of wound response signature genes and *Hopx in vitro* (Figure 2F). Using the aforementioned single cell transcriptomic data from *in vivo* HBCs and the BisqueRNA(Jew et al., 2020) package, we performed decomposition of the *in vitro* bulk transcriptomes to predict the abundances of these HBC subtypes within the bulk transcriptome. Decomposition using the single cell transcriptomic reference revealed a decrease in the predicted proportions of inactivated HBCs after PMA activation, along with an increase in the predicted proportion of activated HBCs (Figure 2G). Collectively, these analyses indicate there is a bidirectional association between the gene expression of *in vitro* and *in vivo* activated HBCs with regards to individual marker genes and activation-associated gene sets.

### Activating in vitro HBCs with PMA induces developmental transcriptional regulatory networks and unlocks bipotent engraftment capacity

Having demonstrated a correspondence between activation by injury *in vivo* and activation by PMA treatment *in vitro,* we examined the transcriptional networks that are differentially regulated as activation proceeds in response to PMA. Previous analyses of the stages of HBC activation *in vivo* have identified transcription factors that regulate aspects of HBC dormancy, activation, and differentiation. As previously mentioned, TP63 is known to regulate HBC dormancy, RELA seems to participate in the activation process, and HOPX is suggested to play a role in the regulation of HBC differentiation. To investigate the roles of these and other transcription factors in the regulation of PMA-activated HBCs, we performed differential expression analysis with DESeq2 to identify differentially expressed transcription factors (n = 249) across the activation timepoints. We then analyzed the sets of TFs and their proposed targets (collectively “regulons”) according to gene co-expression analysis utilizing the R package RTN (version 12.1) (Castro et al., 2015; Corces et al., 2018; M. N. C. Fletcher et al., 2013)(Figure 3A). Our analysis revealed that the 249 transcription factors were associated with 4736 proposed target genes (Figure 3A). The collective expression of the transcription factors and their proposed target genes (regulon activity profiles, RAP) clustered into 5 distinct signatures which changed dynamically across the HBC activation process. The three most prominent regulon clusters corresponded to the vehicle-treated state, early activation (6 HPT), and extended activation (12 HPT).

**Figure 3.**
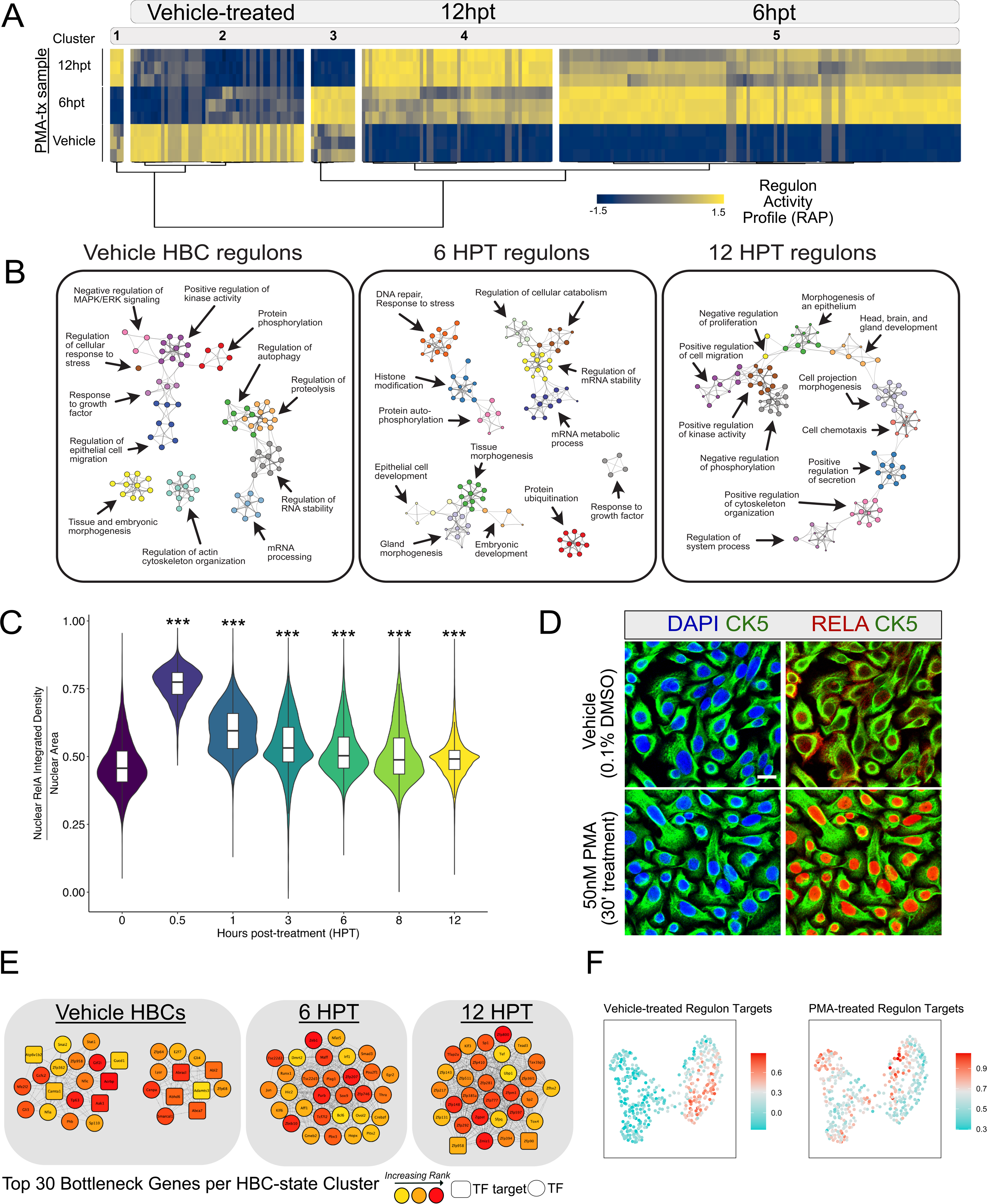
Regulon analysis of PMA-treated HBCs reveals distinct networks underlying activation. A) A heatmap representing the regulon activity profiles exported from RTN. Each line represents an individual regulon (TF + target genes). Clustering is generated using Ward’s minimum variance method with Pearson correlations used as the distance used for clustering. B) Metascape enrichment of regulon clusters, visualized as networks. C) An integrated violin and boxplot representing the Nuclear-Cytoplasmic Index of RelA fluorescence over a 12-h long PMA timecourse (n=3). One-way ANOVA followed by Dunnett’s Test. D) Representative immunofluorescence detailing RelA translocation 30’ after PMA treatment. Scale bar: 20 μm. E) The top 30 bottleneck genes identified per regulon cluster. Upper right legend indicates rank of genes, as well as whether a gene is a transcription factor (circle) or a transcription factor target (square). F) Regulon target gene abundances from Figure 3A mapped onto the single-cell HBC data from Figure 2A for both vehicle-treated and PMA-treated transcription factors. Red indicates high expression and cyan indicates low expression.

We queried each cluster for TFs previously characterized in HBCs. As expected, we identified *Tp63* amongst the resting-state regulons, which suggests a transcriptional network important for the maintenance of inactivated HBCs and serves as an important positive control for this analytic approach. The acute-activation regulons included *RelA.* The extended-activation regulons included *Hopx*(Gadye et al., 2017) suggesting that this cluster may represent gene expression changes important for HBC differentiation. This was supported by ontology analysis of the transcription factors and their target genes: the vehicle-treated regulons were associated with tissue and embryonic morphogenesis, growth factor response, and actin reorganization; the early activation regulons were associated with DNA repair, protein ubiquitination, and morphogenesis suggestive of signaling and metabolic stress; and the extended activation regulons were associated with developmental processes and morphogenesis including cell adhesion and cytoskeletal remodeling (Figure 3B).

Given the *in vivo* findings implicating *RelA* as a crucial effector of HBC activation and the present findings in our study which mapped *RelA* regulon activity to early stages of activation, we sought to determine whether PMA treatment induced *RelA* activity by establishing its nuclear translocation kinetics during the 12-hour long PMA-treatment protocol. Results from this experiment revealed a robust nuclear translocation event taking place just 30 minutes after PMA treatment, which slowly tapered towards baseline-levels nuclear-translocation index (NCI) over the remaining protocol time (Figure 3C, D).

### Topology analyses of regulon communities reveal bottleneck genes relevant for HBC activation

Transcriptional networks, like those identified in the regulon analysis, exist to shape cell states. Our regulon clusters present orchestrated sets of genes that, together, mold a cell’s transcriptional potential for differentiation. To extend this analysis, we were interested in assessing the topological features of our regulon networks. Assessing the topology of biological networks offers the possibility of identifying individual genes whose presence in the network is important for maintaining other biological relationships. We utilized Cytoscape’s (3.8.2) cytoHubba(Chin et al., 2014) plug-in to identify the top 30 bottleneck genes in our regulon clusters (Figure 3C).

Inspection of the top 30 bottleneck genes identified previously validated regulators. As expected, we identified *Tp63* as a bottleneck gene in our vehicle-treated HBC cluster, whose RAP decreases dramatically following PMA activation (Figure 3A and 3C). Given the master regulator role of *Trp63* in HBC activation, its identification as a bottleneck gene suggests that the other 29 genes are relevant for inhibiting activation. Other bottleneck genes found in this group include *Stat1*, *Gli3*, and *Nfe2l2* (Figure 3E, left). Analysis of our 12HPT cluster identified *Hopx* as a top 30 bottleneck gene. The RAP of this cluster steadily increases as a function of PMA-mediated activation (Figure 3A), with their maximal activity taking place at 12HPT, implying that these regulon clusters are important sustaining activation and/or differentiation. Other genes in this group include *Zeb1*, *Jun*, *Sox9,* and *Egr2* (Figure 3E, right). Our 6HPT cluster represents RAPs which maximally increase at 6 HPT but then initiate a return to baseline at 12 HPT (Figure 3A). This suggests that these genes might be functioning analogously to immediate early genes (IEGs), which are rapidly transcribed genes that underlie cellular plasticity(Kim et al., 2018; Minatohara et al., 2016; Okuno, 2011). Genes in this list include *Tox4, Tead3*, *Klf3,* and *Tef* (Figure 3E, middle).

As reported by the authors, transcription factors alone could not distinguish between HBC states in their single-cell RNA seq data se t(Gadye et al., 2017). Consequently, we looked instead at the target genes induced by the transcription factors in our regulon analysis to determine if these could serve as a proxy for the activation process *in vivo*. We found that target genes regulated by vehicle-treated transcription factors were enriched in the partially activated population (Krt5+/Trp63^MED^/Mki67+/Hopx-) (Figure 2A). This result was consisted with our previous report that our primary culture HBC system allows expansion of HBCs that can proliferate but not differentiate(Peterson et al., 2019), and thus are closer to this partially activated population at baseline. In contrast, we found that that our collapsed PMA target genes were the most strongly enriched in the fully activated HBC population (Krt5+/Trp63^none^/Mki67-/Hopx+).

### Transplantation of PMA-activated HBCs into the OE of lesioned host mice

The results from our PMA recovery assay prompted us to consider whether PMA activation resulted in a deterministic commitment to differentiation, or whether the recovery we observed reflected a resistance to activate. To specifically test this question, we carried out transplantation assays of HBCs at the same two stages of activation (6HPT and 12HPT) (Figure 4A). Utilizing an HBC cell line derived from pan-GFP expressing transgenic rats, we transplanted Vehicle-, 6HPT-, and 12HPT HBCs into MeBr-lesioned host rats. After allowing the HBCs to engraft and allowing the OE to regenerate for 12 days after transplantation, we then harvested tissue and quantified the composition of GFP-positive clones (Figure 4A-4B).

**Figure 4.**
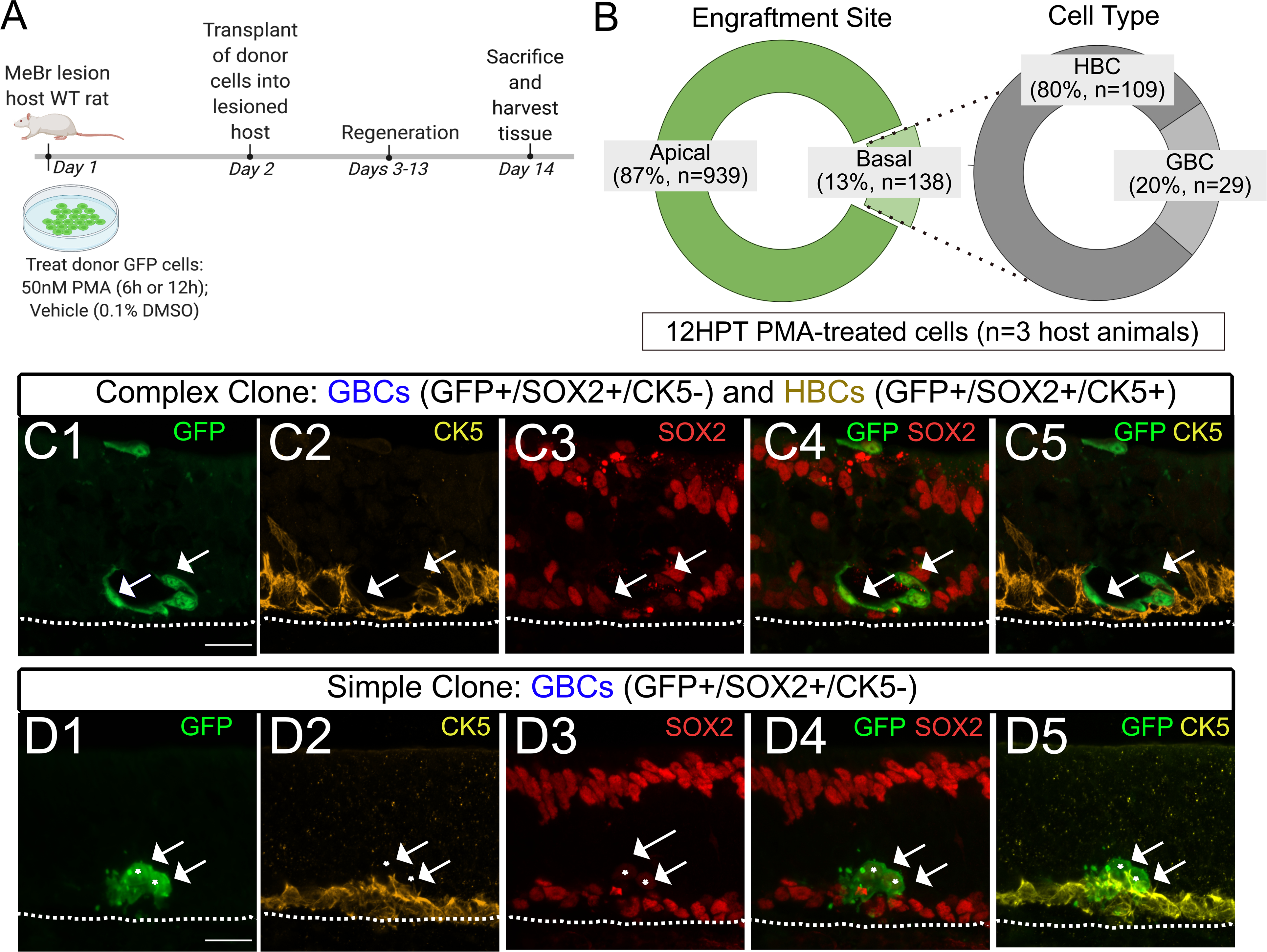
PMA treatment unlocks engraftment and differentiation potential in 12HPT HBCs. A) Schematic illustration of transplantation assay. B) Quantification of 12H PMA-treated GFP+ HBCs (n=1077) based on engraftment location: apical (n=939 total cells) or basal (n=138 total cells). n=3 host rats. Quantification of 12H PMA-treated basal cells (n=138) based on progeny outcome (HBC or GBC). C1-C5) Representative complex clone comprised of HBCs (GFP+/SOX2+/CK5+) and GBCs (GFP+, SOX2+, CK5-). Note representative unengrafted HBC atop the apical cell layer. Scale bar: 20 μm. D1-D5) Representative simple clone comprised of GBCs (GFP+, SOX2+, CK5-). Scale bar: 20 μm.

Vehicle-treated HBCs failed to engraft and differentiate, a result consistent with previous results showing that donor HBCs need to be activated to successfully engraft (Peterson et al., 2019; Schnittke et al., 2015). In stark contrast, we found that 12 hours of PMA-treatment induced a previously undescribed transplant outcome for HBCs wherein HBCs engraft but remain localized on the apical tissue layer. We found that 6HPT failed to engraft at either basal or apical sites, making their transplantation outcome indistinguishable from vehicle-treated cells. The behavior of 12HPT donor HBCs led to two distinct outcomes: first, 87% of GFP+ cells (n=1077 total) were seen resting on the apical layer of the regenerated tissue. These apical GFP+ cells were often found as single cells, but sometimes appeared as doublets. (Figure 4H). The remaining 13% of GFP+ cells (n=138) engrafted in the basal tissue compartment. However, when assessed, most of these cells closely resembled HBCs (80%), while the smaller fraction were at a remove from the basal lamina as would be typical of GBCs (20%) (Figure 4B). To confirm the morphological identification of HBCs vs. GBCs, we immunostained sections containing GFP+ clones for CK5 and SOX2. Cells expressing CK5 and SOX2 were classified as HBCs, while cells expressing SOX2 without CK5 were classified as GBCs. Based on this marker panel, 80% of the engrafted basal cells expressed SOX2(+)/CK5(+), and thus phenotypically HBCs whereas 20% were SOX2(+)/CK5(-) and thus phenotypically GBCs (Figure 4B). Additional features of the engrafted cells are worthy of note. The GFP+ cells tended to form pits near the base of the epithelium, identified by the lack of DAPI+ cells and interruptions in the dense cellularity of the epithelium. Additionally, although most of the clones were simple, we did identify scattered complex clones, some of which contained both SOX2(+)/CK5(+) and SOX2(+)/CK5(-) cells (Figure 4C1-C5).

Collectively, the transcriptional regulatory signatures and *in vivo* engraftment data suggest that treatment of cultured HBCs with PMA for 12 hours unlocks developmental programs sufficient to drive engraftment and forward-differentiation when transplanted into the regenerating OE.

## Discussion

Together, these sets of experiments are the first, to our knowledge, to shed mechanistic insight into the activation behavior of HBCs in the early activation of HBCs, analogous the period immediately after injury. These results have important implications for the study of heterogeneity within the HBC population as a whole and of activation-dormancy transition mechanisms more broadly. Additionally, our results reveal that the biological phenomenon of HBC activation can be successfully modelled *in* vitro. This system can be leveraged to advance future investigations of basic HBC biology and advance the translational potential of cell-based treatments for olfactory dysfunction involving HBCs.

OE regeneration studies have established the first principles governing the major molecular events that arise within HBCs following injury. However, significant efforts must now be placed on articulating the understudied features of HBCs, such as their heterogeneity, plasticity, recovery post-injury, regulation of the TF *Tp63*, and transition across the activation-dormancy-activation axis. For example, as a case study, cell-specific genetic deletion of *Tp63* in an uninjured OE vs. activation by Sus cell ablation produce biased distributions of HBC progeny; OSNs predominate in the former while Sus cells do so in the latter(Herrick et al., 2017)

In this study we show that HBC activation mechanisms, developmental plasticity, and even intercellular heterogeneity can be modelled by PMA treatment with a temporal resolution previously unobtainable. Under this paradigm, we observed a precipitous decline in TP63 occurring between thirty minutes and one hour after activation contemporary with a robust RELA translocation event, thus implicating NFKB signaling in the degradative regulation of TP63 and identifying a precise window of time in which to study TP63 degradation. We focused on two timepoints in the response to the treatment with PMA, specifically early and later stages of activation. At 6 HPT, HBCs begin to engage mechanisms of plasticity as they respond to the metabolic demands of activation. This is evident by their transcriptional regulatory signatures, and their functional bifurcation into cells that either can or cannot recover vehicle-treated TP63 levels following withdrawal of the PMA stimulus. Finally, at 12 HPT, HBCs undergo a second decline in TP63 levels concurrent with HOPX expression, upregulation of developmental and morphogenic transcriptional regulatory signatures, and a unique functional capacity to engraft and forward differentiate *in vivo*. These latter timepoints afford the opportunity to study mechanisms of activation signaling and fate commitment. Results from the PMA recovery assay suggests that PMA-treated HBCs have yet to reach a deterministic critical threshold where TP63 protein synthesis is unrecoverable. This is not surprising, since the culture conditions used to cultivate the HBCs include plated GBCs to differentiate into HBCs themselves(Peterson et al., 2019). Interestingly, regarding reversibility, we find that within this immediate response to PMA treatment, both acute and latently activated HBCs comprise two distinct subpopulations of HBCs, each with varying degrees of TP63 resynthesis potential.

Results from the transplantation assay suggest that the stimulation by PMA was insufficient to mount full differentiation. To expand on this, it was our expectation that PMA could “kickstart” the process and that the post-injury tissue context would still be sufficient to push donor PMA-activated HBCs along this cascade. However, inability to drive large scale regeneration efforts implies that injury signals instead cast HBCs into a plastic state, and that further differentiation requires additional signals that we have not yet been able to replicate completely. In addition, we do not know if the injury process specifies all HBCs to activate in a more or less stochastic manner or whether they are inherently biased toward activating or not prior to injury. For example, in the original demonstration of HBC activation during tissue recovery(Packard et al., 2011), a large proportion of the HBC population showed no evidence of activation, such that the only labeled cells in substantial portions of the epithelium were only HBCs. Of note, FACS analysis of various cell passages showed HBC cultures are relatively homogenous with respect to TP63 and CK5(Peterson et al., 2019). Together, this suggests that TP63 may not be the only factor relevant for the decision of whether to activate or not. However, other studies do suggest alternative explanations for the heterogeneity we see in response to PMA or *in vivo* injury. It has been shown that stem cell differentiation can be a stochastic process and, interestingly, experiments where cells were exposed to the same differentiation stimulus were found to progress through differentiation in an erratic fashion(Stumpf et al., 2017). In this experiment, the authors exposed two different mouse embryonic stem cell lines to a well-established neuronal differentiation protocol and sequenced cells at various time points(Stumpf et al., 2017). While both cell lines eventually “arrived” and their final, intended neuronal state, they found significant variability within and across the cultures. Fundamentally, they found increased cell-to-cell variability while the cells were transitioning from a pluripotent state, which they observe has taken place in other settings(Stumpf et al., 2017). Consequently, it may be the case that HBC activation is also a stochastic process, and that exiting dormancy and moving into activation and differentiation might also be uncoordinated and reflective of this principle. That our primary culture system of HBCs responds heterogeneously to PMA treatment may offer a means of addressing that question. From a clinical perspective this could be useful in dissecting the activation process in substantially more detail.

Our work also presents a pipeline that pairs regulon analysis with topological analyses of these networks to comprehensively characterize and identify regulators for further mechanistic testing. Previous analyses of transcription factor networks have highlighted the notion that the relationship between a transcription factor and its targets provide a better tissue/cell-specific signature than the expression of transcription factors alone(Sonawane et al., 2017). We applied this perspective to our transcriptomic datasets, and to identify novel regulators, used topological analyses prioritize bottleneck genes, which are thought to more likely be essential(Gibney et al., 2013; Yu et al., 2007). Not only did this complementary analysis reveal novel candidates but encompassed candidates that had been previously identified in other experimental paradigms and systems, emphasizing the robustness of this approach. For example, *Nfe2l2*, identified in the dormancy cluster, has been shown to synergistically cooperate with *Tp63* in proliferating keratinocytes to sustain proliferation and promote differentiation(Kurinna et al., 2021), suggesting that *Tp63* likely coordinates HBC dormancy through engagement with other transcription factors.

In summary, we have leveraged our previously established primary *in vitro* HBC culture model (Peterson et al., 2019) to establish a physiologically relevant platform to study HBC activation.

## Experimental procedures

### Resource availability

Raw files, counts, and normalized counts from our bulk-RNA seq can be accessed under GSE215797. Single-cell RNA seq data used in this analysis was accessed from GEO, GSE99251. All raw data, associated code and CellProfiler Pipelines for this manuscript can be accessed at request.

### Animals and breeding

All animals were maintained on *ad libitum* standard rodent chow and water in an AALAC-accredited *vivarium* operating under a 12-hour light/dark cycle. All procedures used on animals were approved by the Committee for the Humane Use of Animals at Tufts University School of Medicine. Eight 10-week-old Sprague Dawley (Taconic NTac:SD) rats were used to generate primary WT HBC lines; these were maintained from a founder breeder pair. Pan-GFP expressing transgenic rats (SD-Tg(ACT-EGFP) CZ-004OOsb strain) were bred and maintained in-house and used to generate pan-GFP HBC lines(Jang et al., 2008; Sasaki et al., 2001).

### Tissue harvest and preparation for cryosectioning

Animals were deeply anesthetized with single, lethal intraperitoneal (IP) injection of triple cocktail of acepromazine (1.25mg/kg), ketamine (37.5mg/kg), and xylazine (7.5mg/kg). Animals were then flushed with an intracardiac perfusion of 10mL of 1X PBS followed by 35mL of ice-cold formalin fixative. The OE was dissected and then incubated with 30mL of formalin for 2 hours at room temperature (RT) and under vacuum. Tissue was then moved to a decalcifying solution of saturated EDTA for at 4C for 48h, before being cryoprotected in 30% sucrose-PBS solution at 4°C for 24 hours. Tissue was then embedded in OCT compound, snap frozen in liquid nitrogen, and stored at −80°C before being sectioned on a Leica cryostat. OE samples were sectioned coronally at 10-20 um of thickness, mounted on Superfrost Plus glass slides (Fisher Scientific, #12-550-150), and stored at −20°C. Rat OE was similarly processed, apart from the perfusion volume which was 200mL of formalin.

### Olfactory epithelium injury assays

Mice were subject to a single intraperitoneal (I.P) injection of methimazole (Sigma-Aldrich, Catalogue - #046KO705) dissolved in 1X sterile PBS (Gibco, Catalogue - #10010-023) at a concentration of 75mg/kg(Håglin et al., 2021; Lin et al., 2017). Seven-to-eight-week-old rats were subjected to MeBr exposure via passive inhalation of 330-345 ppm MeBr mixed with pure air for 6 hours(Schwob et al., 1995).

### Transplantation

P12-P14 rat GFP cells were utilized as donor cells. Host rats were subjected to 330pm-345 ppm MeBr lesion for 6 hours, with the transplants taking place 20 hours after the termination of the MeBr exposure. Donor cells were treated with 50nM PMA (6 hours or 12 hours) or 0.1% DMSO for vehicle control. Cells were grown to 90% confluency on a 100mm poly-d-lysine/laminin coated dishes, in triplicate. 8-to-9-week-old littermate male rats were used as host animals. Upon transplantation time, cells were washed 3X with PBS and treated with Accutase at RT until detachment. Upon detachment, cells were spun down and resuspended in 150uL DMEM/F12 and kept on ice.

The anterior neck of the host animals was shaved to permit access to the trachea. The skin was sanitized and disinfected with prep pads pre-soaked with 70% isopropyl alcohol (Covidien, Catalogue #57520) and then iodine. A tracheotomy was performed, and the palate was raised with a 3 cm piece of PE-100 tubing to close the nasopharyngeal passage and prevent flooding the lungs with the DMEM/F12 cell mixture. Cells were gently resuspended and aspirated into a 1mL plastic syringe (Fisher Scientific, Catalogue #14955456). Cells were then slowly infused into one naris until liquid could be seen starting to exit the other naris. The tracheotomy was then sutured, rats were injected with 2mL of saline, and rats were placed onto a warm heat pad singly housed and allowed to recover overnight.

### Immunohistochemistry

Slides were washed with 1X PBS before baking on a plate warmer (65°C) for one hour. All pre-treatments and staining conditions took place in a humidified chamber. Tissue sections were then pretreated with a 5-minute incubation of 3% H_2_O_2_ in MeOH and 10 minutes of antigen retrieval in a commercial food steamer (incubated with 0.01M pH 6.0 citrate buffer). Sections were then subjected to blocking with block buffer (10% normal donkey serum, 5% nonfat dry milk, 4% BSA, 0.1% Tritonx-100 in 1X PBS) for one 1 hour at RT. Primary antibodies were prepared in block buffer and incubated for either 1 hour at RT or 16 hours overnight at 4°C. Antibody concentrations can be found in Supplemental Table 1.

### *In vitro* model of horizontal basal cells (HBCs) and PMA activation

HBCs were cultured using our established primary culture HBC system(Peterson et al., 2019) with slight modifications to the antibiotic regimen. Instead of utilizing Penicillin-Streptomycin, we used Primocin (100 μg/ml final concentration) and anti-mycotic/anti-biotic (1X final concentration, Gibco Catalogue #15240062). For PMA-mediated activation, HBCs were treated with 50nM PMA (Sigma Aldrich Catalogue #P8139) resuspended in their maintenance media for up to 12 hours.

### Immunocytochemistry

Upon the end of indicated treatment lengths and times, cells were washed with 1X PBS three times for 5 minutes. Cells were then fixed with 10% formalin for 15 minutes at RT. Upon three 1X PBS, 5-minute washes, cells were permeabilized with ice-cold methanol on ice. Cells were then blocked for 1hr at RT utilizing the block buffer (10% normal donkey serum, 5% nonfat dry milk, 4% BSA, 0.1% Tritonx-100 in 1X PBS). Primary antibodies were prepared in block buffer and incubated for either 1 hour at RT or 16 hours overnight at 4°C. Antibody concentrations can be found in Supplemental Table 1.

### Confocal microscopy

For *in vitro* imaging experiments, HBCs were grown and treated on poly-d-lysine/laminin coated glass coverslips or glass-bottom chamber slides. All samples, either tissue sections or *in vitro* HBCs, were imagined on a Zeiss LSM800 confocal microscope utilizing the multi-track mode. For the PMA-induced TP63 loss time-course, the highest TP63 fluorescence Z-level was manually selected for each sample before creating a 5×5 mosaic imaging grid. All other *in vitro* experiments were imaged by creating a 2×2 mosaic grid with a multi-step (5-8 step) Z-stack, before being maximally projected in Fiji prior to analysis. All representative image samples underwent a Z-stack imaging approach utilizing Zeiss software recommendations before being maximally projected in Fiji. Images were background corrected utilizing the Rolling Ball Algorithm in ImageJ. Representative images were prepared in Fiji before being assembled in Affinity Designer.

### CellProfiler for quantitative image analysis

CellProfiler was used to quantify fluorescent images following pre-processing of images as described in these methods. Pipelines are available at request.

### Bulk RNA transcriptomic sequencing

Following indicated treatment lengths, cultured HBCs were washed 3x with PBS and incubated with Accutase at RT for 15 minutes. Suspended cells were spun and washed with 1X PBS before being flash frozen in liquid nitrogen and kept at −80°C overnight. Total RNA was harvested from frozen cell pellets utilizing the PureLink RNA Mini Kit per manufacturer’s instructions. Residual DNA was removed utilizing the PureLink DNAse Set, according to manufacturer’s instructions, before being subjected to NanoDrop analysis. RNA QC, cDNA synthesis, and library preparation was carried out by Novogene Corporation Inc. Samples underwent robust quality control (QC) assessment by Novogene Corporation, which included an assessment of RNA quality (RNA Integrity Number [RIN], all samples ≥ 9.7), cDNA library QC, which included library quantification and insert size assessment, as well as sequencing quality control Samples were loaded onto an Illumina NovaSeq 6000 sequencer and samples were underwent paired end 150bp sequencing (PE150). Following demultiplexing, samples were delivered via FTP.

### Preprocessing and quality control of bulk RNA-sequencing samples and differential gene expression (DEG)

Samples were aligned to the rat genome (rn6, version 2.7.4a) utilizing STAR(Dobin et al., 2013) (version 2.7.5b) with the following settings: “—quantMode GeneCounts”. Samples were normalized using by using the estimateSIzeFactors function from DEseq2(Love et al., 2014). Differential gene expression (DGE) analysis was carried out utilizing the ‘results’ function of DESeq2. The alpha was set to 0.05.

### Topological analyses of dynamically changing regulon networks and bottleneck gene identification

Regulon analysis was carried out with the R package RTN (v. 2.12.1). A list of rat transcription factors (TFs) was accessed from the Animal Transcription Factor Database(Hu et al., 2019) (v. 3.0), cross-referenced with our normalized matrix counts for each sample, and 249 TFs were inputted. Standard package recommendations were followed, except for the deletion of the bootstrapping step due to limited sample size. Identified regulon activity profiles (RAPs) were exported directly into Cytoscape(Shannon et al., 2003) (v. 3.8.2) utilizing the R package RCy3 (v. 2.8.1) for further analysis. The five RAP clusters underwent further topological analysis to identify bottleneck genes utilizing the Cytoscape (v. 3.0) package CytoHubba(Chin et al., 2014) (v. 0.1).

### Statistics and data visualization

All statistical analyses were carried out in R/R-Studio (R version 4.0.2, R-Studio version 1.3.1056). Visualization was performed with the package ggplot2 (version 3.3.3), unless otherwise indicated in the figure legends. For groups with multiple comparisons, a one-way ANOVA followed by Dunnett’s Test was calculated, relative to untreated or vehicle condition. Information about sample sizes and other statistical tests can be found in their respective figure legends. Sample size information for *in vitro* can be found under Supplemental Information S1. Significance levels are demarcated by asterisks, which are as follows: *p < 0.05, **p <0.01, ***p <0.001.

## Author contributions

C.B.C., conceptualization; C.B.C, data curation; C.B.C, M.J.Z, J.D.L, formal analysis; J.E.S, funding acquisition; C.B.C, J.D.L, W.J., M.J.Z., investigation; C.B.C, J.D.L., and M.J.Z,., methodology; J.E.S, supervision; C.B.C, writing – original draft; C.B.C., J.D.L., M.J.Z., J.E.S: writing – review & editing., All authors contributed to the article and approved the submitted version.

## Acknowledgements

The authors wish to thank Po Tse Kwok for her outstanding technical contributions to this project. The authors also wish to thank Drs. Greg Bashaw and Brian Lin for their critical feedback on this manuscript.

---

**<u>Table S1</u>** Primary antibodies used in this study, their source, and their concentration.

<u>Supplemental Information S1</u>

Sample size information for *in vitro* experiments.

**Supplemental Figure S1.**
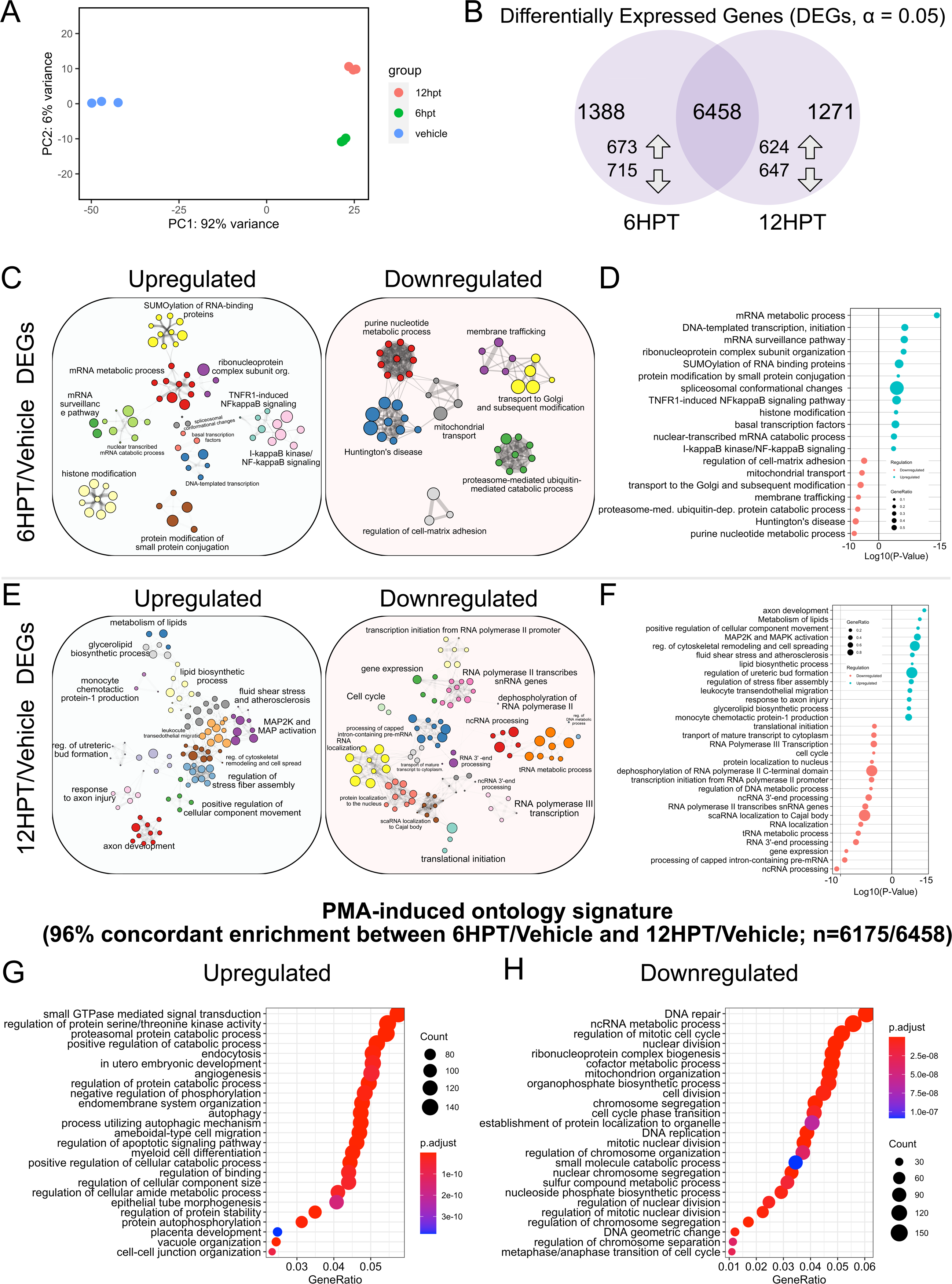
Transcriptomic characterization of PMA-treated HBCs. A) Principal component analysis (PCA) plot of 12h-, 6h- and vehicle-PMA treated HBCs (n=3). B) Venn diagram of differentially expressed genes (DEGs) of 12h- and 6h-PMA treated HBCs relative to vehicle. DEGs were identified with a false discovery rate (FDR) adjusted p-value of < 0.05. C) Metascape enrichment of 6HPT/Vehicle DEGs genes, visualized as ontology networks D) Accompanying ontology dotplot of significantly regulated ontological categories from (C) E) Metascape enrichment of 6HPT/Vehicle DEGs genes, visualized as ontology networks F) Accompanying ontology dotplot of significantly regulated ontological categories from (E) G) ClusterProfiler dotplot of GO Biological Process ontology for shared, upregulated 6HPT and 12HPT genes, relative to vehicle. H) ClusterProfiler dotplot of GO Biological Process ontology for shared, downregulated 6HPT and 12HPT genes, relative to vehicle. For (D) and (F) Significance testing determined by Metascape. For (G) and (H) FDR adjusted p-value < 0.05 and q-value < 0.10. Ontology categories were simplified utilizing the Wang method, with a similarity cutoff of 0.7.

## References Cited

Carr, V. M., Menco, B. Ph. M., Yankova, M. P., Morimoto, R. I., & Farbman, A. I. (2001). Odorants as cell-type specific activators of a heat shock response in the rat olfactory mucosa. The Journal of Comparative Neurology, 432(4), 425–439. 10.1002/cne.1112

Carter, L. A., MacDonald, J. L., & Roskams, A. J. (2004). Olfactory horizontal basal cells demonstrate a conserved multipotent progenitor phenotype. The Journal of NeurosciencelJ: The Official Journal of the Society for Neuroscience, 24(25), 5670–5683. 10.1523/JNEUROSCI.0330-04.2004

Castro, M. A. A., de Santiago, I., Campbell, T. M., Vaughn, C., Hickey, T. E., Ross, E., Tilley, W. D., Markowetz, F., Ponder, B. A. J., & Meyer, K. B. (2015). Regulators of genetic risk of breast cancer identified by integrative network analysis. Nature Genetics, 48(1), 12–21. 10.1038/ng.3458

Chen, M., Reed, R. R., & Lane, A. P. (2017). Acute inflammation regulates neuroregeneration through the NF-κB pathway in olfactory epithelium. Proceedings of the National Academy of Sciences of the United States of America, 114(30), 8089–8094. 10.1073/pnas.1620664114

Chin, C.-H., Chen, S.-H., Wu, H.-H., Ho, C.-W., Ko, M.-T., & Lin, C.-Y. (2014). cytoHubba: identifying hub objects and sub-networks from complex interactome. BMC Systems Biology, 8 Suppl 4(Suppl 4), S11–S11. 10.1186/1752-0509-8-S4-S11

Corces, M. R., Granja, J. M., Shams, S., Louie, B. H., Seoane, J. A., Zhou, W., Silva, T. C., Groeneveld, C., Wong, C. K., Cho, S. W., Satpathy, A. T., Mumbach, M. R., Hoadley, K. A., Robertson, A. G., Sheffield, N. C., Felau, I., Castro, M. A. A., Berman, B. P., Staudt, L. M., … Zhu, J. (2018). The chromatin accessibility landscape of primary human cancers. Science, 362(6413), eaav1898. 10.1126/science.aav1898

Dobin, A., Davis, C. A., Schlesinger, F., Drenkow, J., Zaleski, C., Jha, S., Batut, P., Chaisson, M., & Gingeras, T. R. (2013). STAR: ultrafast universal RNA-seq aligner. *Bioinformatics (Oxford*, England), 29(1), 15–21. 10.1093/bioinformatics/bts635

Fletcher, M. N. C., Castro, M. A. A., Wang, X., de Santiago, I., O’Reilly, M., Chin, S. F., Rueda, O. M., Caldas, C., Ponder, B. A. J., Markowetz, F., & Meyer, K. B. (2013). Master regulators of FGFR2 signalling and breast cancer risk. Nature Communications, 4. 10.1038/ncomms3464

Fletcher, R. B., Das, D., Gadye, L., Street, K. N., Baudhuin, A., Wagner, A., Cole, M. B., Flores, Q., Choi, Y. G., Yosef, N., Purdom, E., Dudoit, S., Risso, D., & Ngai, J. (2017). Deconstructing Olfactory Stem Cell Trajectories at Single-Cell Resolution. Cell Stem Cell, 20(6), 817–830.e8. 10.1016/j.stem.2017.04.003

Gadye, L., Das, D., Sanchez, M. A., Street, K., Baudhuin, A., Wagner, A., Cole, M. B., Choi, Y. G., Yosef, N., Purdom, E., Dudoit, S., Risso, D., Ngai, J., & Fletcher, R. B. (2017). Injury Activates Transient Olfactory Stem Cell States with Diverse Lineage Capacities. Cell Stem Cell, 21(6), 775–790.e9. 10.1016/j.stem.2017.10.014

Gibney, P. A., Lu, C., Caudy, A. A., Hess, D. C., & Botstein, D. (2013). Yeast metabolic and signaling genes are required for heat-shock survival and have little overlap with the heat-induced genes. Proceedings of the National Academy of Sciences of the United States of America, 110(46), E4393–E4402. 10.1073/pnas.1318100110

Håglin, S., Bohm, S., & Berghard, A. (2021). Single or repeated ablation of mouse olfactory epithelium by methimazole. Bio-Protocol, 11(8). 10.21769/BioProtoc.3983

Herrick, D. B., Lin, B., Peterson, J., Schnittke, N., & Schwob, J. E. (2017). Notch1 maintains dormancy of olfactory horizontal basal cells, a reserve neural stem cell. Proceedings of the National Academy of Sciences of the United States of America, 114(28), E5589–E5598. 10.1073/pnas.1701333114

Holden, N. S., Squires, P. E., Kaur, M., Bland, R., Jones, C. E., & Newton, R. (2008). Phorbol ester-stimulated NF-κB-dependent transcription: Roles for isoforms of novel protein kinase C. Cellular Signalling, 20(7), 1338–1348. 10.1016/j.cellsig.2008.03.001

Hu, H., Miao, Y.-R., Jia, L.-H., Yu, Q.-Y., Zhang, Q., & Guo, A.-Y. (2019). AnimalTFDB 3.0: a comprehensive resource for annotation and prediction of animal transcription factors. Nucleic Acids Research, 47(D1), D33–D38. 10.1093/nar/gky822

Jang, W., Lambropoulos, J., Woo, J. K., Peluso, C. E., & Schwob, J. E. (2008). Maintaining epitheliopoietic potency when culturing olfactory progenitors. Experimental Neurology, 214(1), 25–36. 10.1016/j.expneurol.2008.07.012

Jang, W., Youngentob, S. L., & Schwob, J. E. (2003). Globose basal cells are required for reconstitution of olfactory epithelium after methyl bromide lesion. The Journal of Comparative Neurology, 460(1), 123–140. 10.1002/cne.10642

Jew, B., Alvarez, M., Rahmani, E., Miao, Z., Ko, A., Garske, K. M., Sul, J. H., Pietiläinen, K. H., Pajukanta, P., & Halperin, E. (2020). Accurate estimation of cell composition in bulk expression through robust integration of single-cell information. Nature Communications, 11(1). 10.1038/s41467-020-15816-6

Kim, S., Kim, H., & Um, J. W. (2018). Synapse development organized by neuronal activity-regulated immediate-early genes. In Experimental and Molecular Medicine (Vol. 50, Issue 4). Nature Publishing Group. 10.1038/s12276-018-0025-1

Kurinna, S., Seltmann, K., Bachmann, A. L., Schwendimann, A., Thiagarajan, L., Hennig, P., Beer, H. D., Mollo, M. R., Missero, C., & Werner, S. (2021). Interaction of the NRF2 and p63 transcription factors promotes keratinocyte proliferation in the epidermis. Nucleic Acids Research, 49(7), 3748–3763. 10.1093/nar/gkab167

Lin, B., Coleman, J. H., Peterson, J. N., Zunitch, M. J., Jang, W., Herrick, D. B., & Schwob, J. E. (2017). Injury Induces Endogenous Reprogramming and Dedifferentiation of Neuronal Progenitors to Multipotency. Cell Stem Cell, 21(6), 761–774.e5. 10.1016/j.stem.2017.09.008

Love, M. I., Huber, W., & Anders, S. (2014). Moderated estimation of fold change and dispersion for RNA-seq data with DESeq2. Genome Biology, 15(12), 550. 10.1186/s13059-014-0550-8

McQuin, C., Goodman, A., Chernyshev, V., Kamentsky, L., Cimini, B. A., Karhohs, K. W., Doan, M., Ding, L., Rafelski, S. M., Thirstrup, D., Wiegraebe, W., Singh, S., Becker, T., Caicedo, J. C., & Carpenter, A. E. (2018). CellProfiler 3.0: Next-generation image processing for biology. PLoS Biology, 16(7), e2005970– e2005970. 10.1371/journal.pbio.2005970

Minatohara, K., Akiyoshi, M., & Okuno, H. (2016). Role of Immediate-Early Genes in Synaptic Plasticity and Neuronal Ensembles Underlying the Memory Trace. Frontiers in Molecular Neuroscience, 8, 78. 10.3389/fnmol.2015.00078

Okuno, H. (2011). Regulation and function of immediate-early genes in the brain: Beyond neuronal activity markers. Neuroscience Research, 69(3), 175–186. 10.1016/j.neures.2010.12.007

Packard, A., Schnittke, N., Romano, R. A., Sinha, S., & Schwob, J. E. (2011). ΔNp63 regulates stem cell dynamics in the Mammalian olfactory epithelium. Journal of Neuroscience, 31(24), 8748–8759. 10.1523/JNEUROSCI.0681-11.2011

Peterson, J., Lin, B., Barrios-Camacho, C. M., Herrick, D. B., Holbrook, E. H., Jang, W., Coleman, J. H., & Schwob, J. E. (2019). Activating a Reserve Neural Stem Cell Population In Vitro Enables Engraftment and Multipotency after Transplantation. Stem Cell Reports, 12(4), 680–695. 10.1016/j.stemcr.2019.02.014

Sasaki, M., Honmou, O., Akiyama, Y., Uede, T., Hashi, K., & Kocsis, J. D. (2001). Transplantation of an acutely isolated bone marrow fraction repairs demyelinated adult rat spinal cord axons. GLIA, 35(1), 26–34. 10.1002/glia.1067

Schnittke, N., Herrick, D. B., Lin, B., Peterson, J., Coleman, J. H., Packard, A. I., Jang, W., & Schwob, J. E. (2015). Transcription factor p63 controls the reserve status but not the stemness of horizontal basal cells in the olfactory epithelium. Proceedings of the National Academy of Sciences of the United States of America, 112(36), E5068–E5077. 10.1073/pnas.1512272112

Schwob, J. E. (2005). Restoring Olfaction: A View from the Olfactory Epithelium. Chemical Senses, 30(Supplement 1), i131–i132. 10.1093/chemse/bjh149

Schwob, J. E., Jang, W., Holbrook, E. H., Lin, B., Herrick, D. B., Peterson, J. N., & Hewitt Coleman, J. (2017). Stem and progenitor cells of the mammalian olfactory epithelium: Taking poietic license. The Journal of Comparative Neurology, 525(4), 1034–1054. 10.1002/cne.24105

Schwob, J. E., Youngentob, S. L., & Mezza, R. C. (1995). Reconstitution of the rat olfactory epithelium after methyl bromide-induced lesion. The Journal of Comparative Neurology, 359(1), 15–37. 10.1002/cne.903590103

Shannon, P., Markiel, A., Ozier, O., Baliga, N. S., Wang, J. T., Ramage, D., Amin, N., Schwikowski, B., & Ideker, T. (2003). Cytoscape: a software environment for integrated models of biomolecular interaction networks. Genome Research, 13(11), 2498–2504. 10.1101/gr.1239303

Sonawane, A. R., Platig, J., Fagny, M., Chen, C.-Y., Paulson, J. N., Lopes-Ramos, C. M., DeMeo, D. L., Quackenbush, J., Glass, K., & Kuijjer, M. L. (2017). Understanding Tissue-Specific Gene Regulation. Cell Reports, 21(4), 1077–1088. 10.1016/j.celrep.2017.10.001

Stumpf, P. S., Smith, R. C. G., Lenz, M., Schuppert, A., Müller, F.-J., Babtie, A., Chan, T. E., Stumpf, M. P. H., Please, C. P., Howison, S. D., Arai, F., & MacArthur, B. D. (2017). Stem Cell Differentiation as a Non-Markov Stochastic Process. Cell Systems, 5(3), 268–282.e7. 10.1016/j.cels.2017.08.009

Tirosh, I., Izar, B., Prakadan, S. M., Wadsworth, M. H., Treacy, D., Trombetta, J. J., Rotem, A., Rodman, C., Lian, C., Murphy, G., Fallahi-Sichani, M., Dutton-Regester, K., Lin, J. R., Cohen, O., Shah, P., Lu, D., Genshaft, A. S., Hughes, T. K., Ziegler, C. G. K., … Garraway, L. A. (2016). Dissecting the multicellular ecosystem of metastatic melanoma by single-cell RNA-seq. Science, 352(6282), 189–196. 10.1126/science.aad0501

Yu, H., Kim, P. M., Sprecher, E., Trifonov, V., & Gerstein, M. (2007). The importance of bottlenecks in protein networks: correlation with gene essentiality and expression dynamics. PLoS Computational Biology, 3(4), e59–e59. 10.1371/journal.pcbi.0030059

Zhou, Y., Zhou, B., Pache, L., Chang, M., Khodabakhshi, A. H., Tanaseichuk, O., Benner, C., & Chanda, S. K. (2019). Metascape provides a biologist-oriented resource for the analysis of systems-level datasets. Nature Communications, 10(1), 1523. 10.1038/s41467-019-09234-6

