## Supplemental Table 1 for "An *in vitro* model of acute horizontal basal cell activation reveals dynamic gene regulatory networks underlying the acute activation phase"

| Target | Source and vendor | Concentration |
|---|---|---|
| Ms α-P63 | ATCC | 1:100 |
| Ch α-KERATIN 5 | Biolegend (#905904) | 1:500 |
| Rb α-KI67 | Novus Biologicals (#11089717F) | 1:250 |
| Rb α-HOPX | Proteintech (11419-1-AP) | 1:250 |
| Rb α-RelA | CST (D14E12) | 1:250 |

Table S1: Primary antibodies, their source, and concentration
